## supplemental figures s1-S12 for "Merging Multi-OMICs with Proteome Integral Solubility Alteration Unveils Antibiotic Mode of Action": Fig S1 - S12.pdf

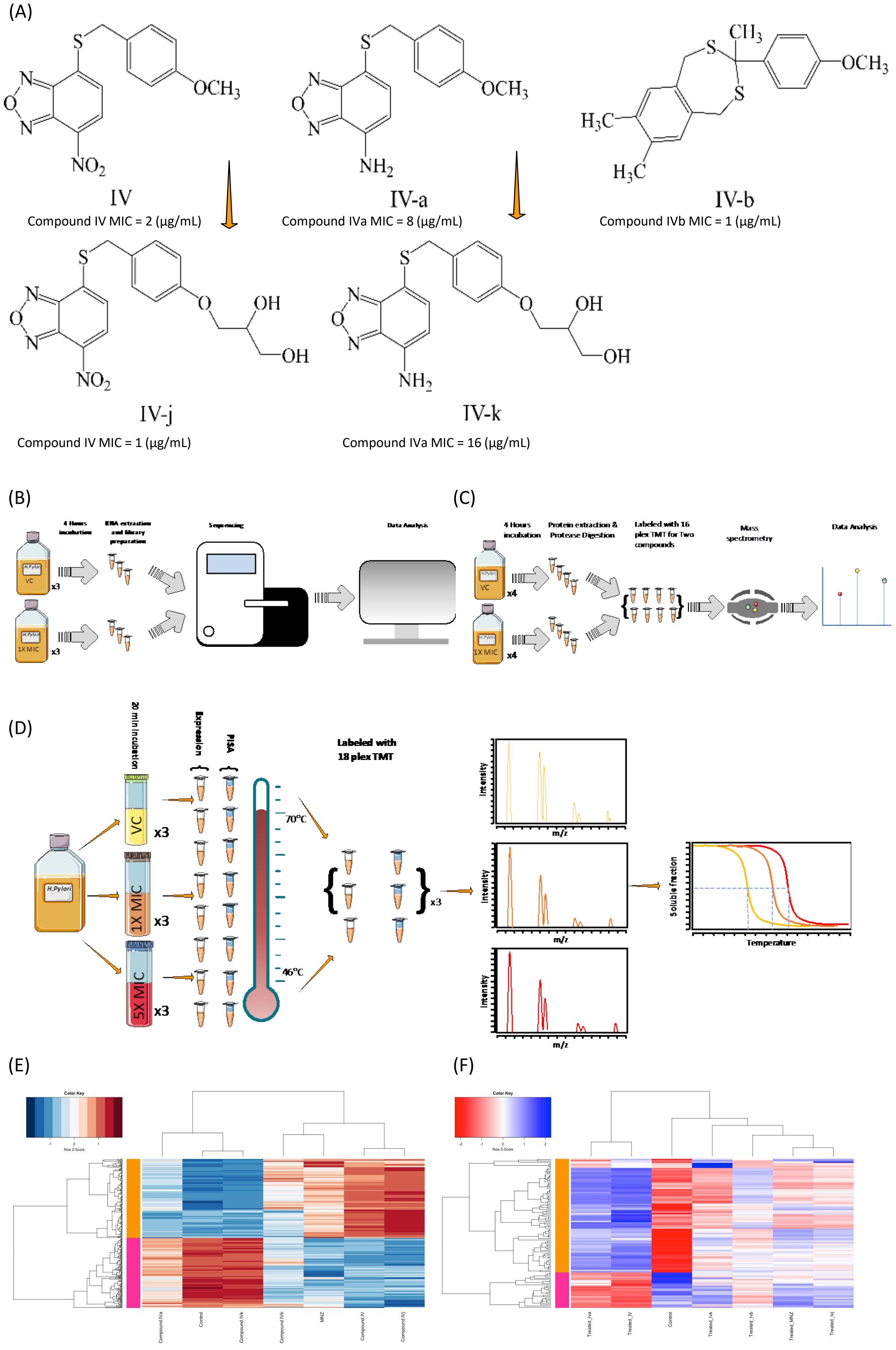
(A)
IV
Compound IV MIC = 2 (µg/mL)
IV-a
Compound IVa MIC = 8 (µg/mL)
IV-b
Compound IVb MIC = 1 (µg/mL)
IV-j
Compound IV MIC = 1 (µg/mL)
IV-k
Compound IVa MIC = 16 (µg/mL)
(B)
Sequencing
Data Analysis
VC
1X MIC
x3
(C)
4 Hours incubation
Protein extraction & Protease Digestion
Labeled with 16 plex TMT for Two compounds
Mass spectrometry
Data Analysis
VC
1X MIC
x4
(D)
20 min incubation
Expression
PISA
Labeled with 18 plex TMT
H.Pylori
VC
1X MIC
5X MIC
x3
70°C
46°C
Intensity
m/z
Soluble fraction
Temperature
(E)
Color Key
Row Z-Score
Compound IVa
Control
Compound IVk
Compound IVb
MNZ
Compound IV
Compound IVj
(F)
Color Key
Row Z-Score
Treated_Iva
Treated_IV
Control
Treated_Ivk
Treated_Ivb
Treated_MNZ
Treated_Ivj

**Fig. S1 Compounds used in the assay and the study design.**

**(A)** The compounds used in this assay were selected from a high-throughput screening that aimed to identify flavodoxin binders from a diverse chemical library of 10,000 compounds. Through several rounds of chemical variation and efficacy testing, a family of novel nitrobenzoxadiazol-based antimicrobials emerged, with compound IV as the lead compound. Compound IVj and IVk are more soluble derivatives of compound IV, and IVa is a derivative of compound IV with an amine functionality. The compounds bearing an amine functionality are narrow-spectrum antimicrobials that exhibit high specificity against H. pylori and potentially other gastric Helicobacter species but not enterohepatic Helicobacter species. The two compounds tested that contain a nitro functionality show extended-spectrum activity against Gram-positive bacteria, the Helicobacter genus, and C. jejuni. These extended-spectrum antimicrobials may have potential for novel therapies against Gram-positive bacteria, while the narrow-spectrum ones may be useful specifically against H. pylori. **(B)** Transcriptomics Study Design for Drug Testing: Provide a description of the transcriptomics study design for drug testing. **(C)** Proteomics Assay Study Design for Drug Testing: Provide a description of the proteomics assay study design for drug testing. **(D)** Proteome Integral Solubility Alteration (PISA) Assay Study Design for Determination of Intracellular Targets: Provide a description of the Proteome Integral Solubility Alteration (PISA) assay study design for determining intracellular targets. **(E)** The heatmap displays 771 differentially expressed genes, and hierarchical clustering is performed based on different compounds. **(F)** The heatmap represents 113 differentially expressed proteins, and hierarchical clustering is conducted based on different compounds.

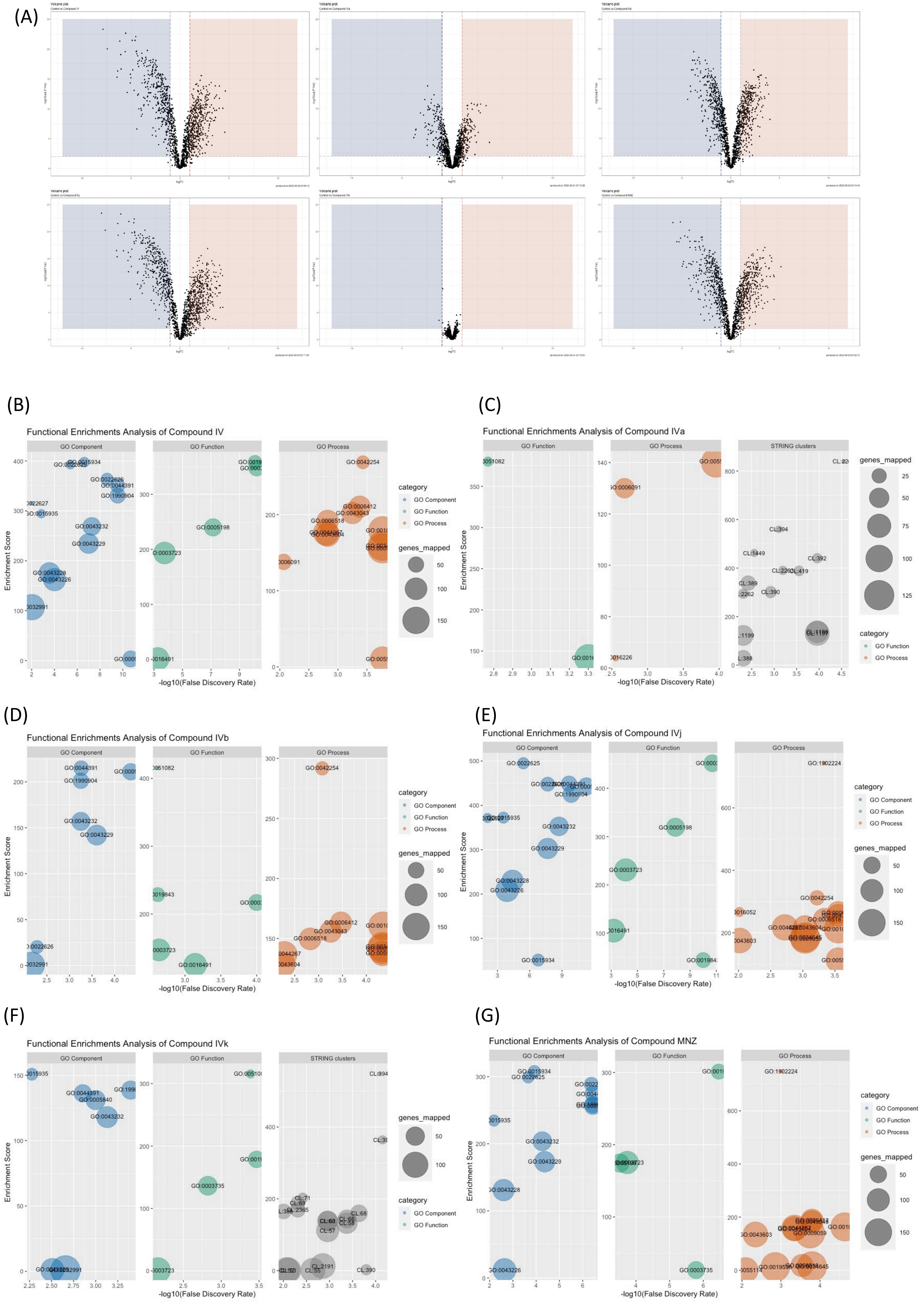

(A)
(B)
Functional Enrichments Analysis of Compound IV
GO Component
GO Function
GO Process
Enrichment Score
-log10(False Discovery Rate)
category
genes_mapped
(C)
Functional Enrichments Analysis of Compound IVa
STRING clusters
(D)
Functional Enrichments Analysis of Compound IVb
(E)
Functional Enrichments Analysis of Compound IVj
(F)
Functional Enrichments Analysis of Compound IVk
(G)
Functional Enrichments Analysis of Compound MNZ

**Fig. S2 Gene ontology analysis of Compound IV and its derivatives**

**(A)** Volcano plot: Displays the differentially expressed genes identified from the transcriptomics assay. **(B)** Distribution of gene ontology components associated with Compound IV: Shows the distribution of gene ontology terms associated with the biological processes, molecular functions, and cellular components influenced by Compound IV. **(C)** Distribution of gene ontology components and String cluster associated with Compound IVa: Presents the distribution of gene ontology terms and String cluster analysis results associated with Compound IVa. **(D)** Distribution of gene ontology components associated with Compound IVb: Illustrates the distribution of gene ontology terms associated with the biological processes, molecular functions, and cellular components influenced by Compound IVb. **(E)** Distribution of gene ontology components associated with Compound IVj: Shows the distribution of gene ontology terms associated with the biological processes, molecular functions, and cellular components influenced by Compound IVj. **(F)** Distribution of gene ontology components and String cluster associated with Compound IVk: Presents the distribution of gene ontology terms and String cluster analysis results associated with Compound IVk, which is a more soluble derivative of Compound IVa. **(G)** Distribution of gene ontology components associated with Compound MNZ: Depicts the distribution of gene ontology terms associated with the biological processes, molecular functions, and cellular components influenced by Compound MNZ.

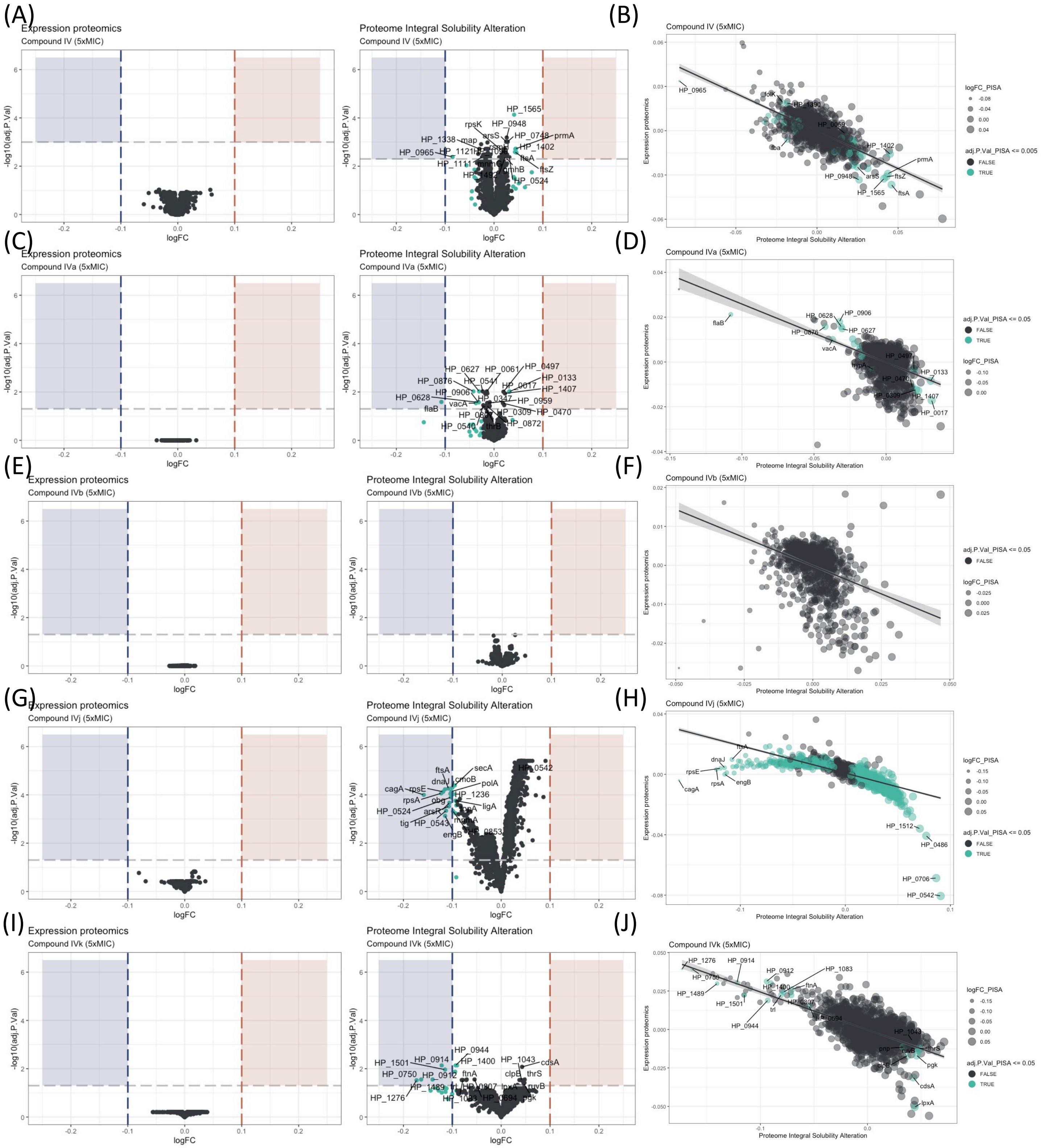

(A)
Expression proteomics
Compound IV (5xMIC)
Proteome Integral Solubility Alteration
Compound IV (5xMIC)
-log10(adj.P.Val)
logFC
(B)
Compound IV (5xMIC)
Expression proteomics
Proteome Integral Solubility Alteration
logFC_PISA
adj.P.Val_PISA <= 0.005
FALSE
TRUE
(C)
Expression proteomics
Compound IVa (5xMIC)
Proteome Integral Solubility Alteration
Compound IVa (5xMIC)
(D)
Compound IVa (5xMIC)
adj.P.Val_PISA <= 0.05
logFC_PISA
(E)
Expression proteomics
Compound IVb (5xMIC)
Proteome Integral Solubility Alteration
Compound IVb (5xMIC)
(F)
Compound IVb (5xMIC)
(G)
Expression proteomics
Compound IVj (5xMIC)
Proteome Integral Solubility Alteration
Compound IVj (5xMIC)
(H)
Compound IVj (5xMIC)
(I)
Expression proteomics
Compound IVk (5xMIC)
Proteome Integral Solubility Alteration
Compound IVk (5xMIC)
(J)
Compound IVk (5xMIC)

**Fig. S3 Proteome Integral Solubility Alteration coupled with expression proteomics (PISA-Express) analysis for different compounds at 5x of MIC.**

**(A)** Changes in expression proteomics data and Proteome Integral Solubility Alteration results in five times of MIC concentrations of Compound IV. **(B)** Comparison of the expression proteomics data with Proteome Integral Solubility Alteration demonstrates significant alterations (teal color) in protein solubility for Compound IV. **(C)** Changes in expression proteomics data and Proteome Integral Solubility Alteration results in five times of MIC concentrations of Compound IVa. **(D)** Comparison of the expression proteomics data with Proteome Integral Solubility Alteration demonstrates significant alterations (teal color) in protein solubility for Compound IVa. **(E)** Compound IVb does not revile any Changes in expression proteomics data and Proteome Integral Solubility Alteration results in five times of MIC concentrations of Compound IVb. **(F)** Comparison of the expression proteomics data and Proteome Integral Solubility Alteration results with five times of MIC concentrations of Compound IVb showing a negative correlation but does not revile any significant changes. **(G)** Changes in expression proteomics data and Proteome Integral Solubility Alteration results in five times of MIC concentrations of Compound IVj. **(H)** Comparison of the expression proteomics data with Proteome Integral Solubility Alteration demonstrates significant alterations (teal color) in protein solubility for Compound IVj. **(I)** Changes in expression proteomics data and Proteome Integral Solubility Alteration results in five times of MIC concentrations of Compound IVk. **(J)** Comparison of the expression proteomics data with Proteome Integral Solubility Alteration demonstrates significant alterations (teal color) in protein solubility for Compound IVk.

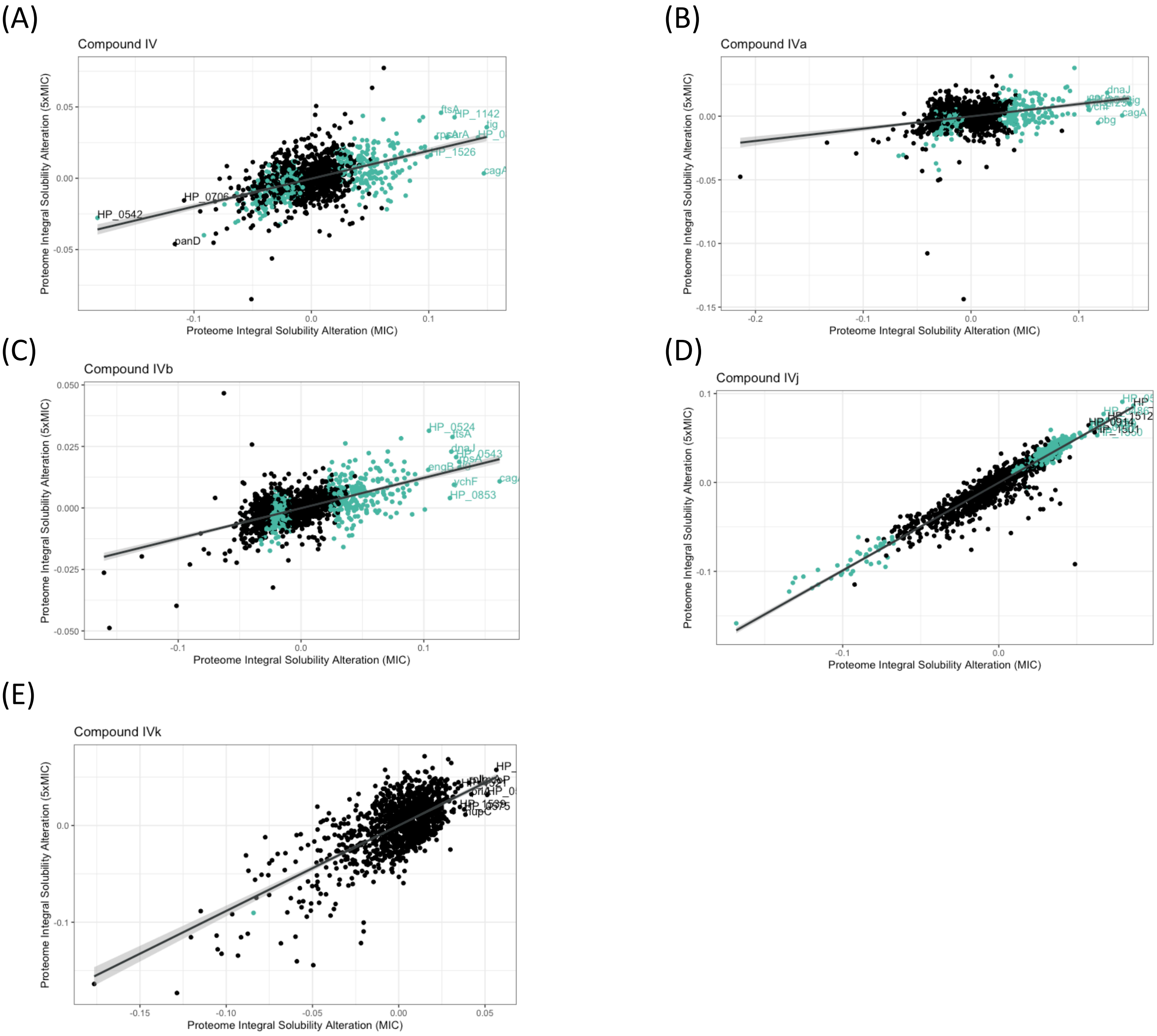
(A)
Compound IV
Proteome Integral Solubility Alteration (5xMIC)
Proteome Integral Solubility Alteration (MIC)
ftsA
HP_1142
lig
HP_1526
cagA
HP_0542
HP_0706
panD
(B)
Compound IVa
Proteome Integral Solubility Alteration (5xMIC)
Proteome Integral Solubility Alteration (MIC)
dnaJ
cagA
obg
(C)
Compound IVb
Proteome Integral Solubility Alteration (5xMIC)
Proteome Integral Solubility Alteration (MIC)
HP_0524
ftsA
HP_0543
ychF
HP_0853
(D)
Compound IVj
Proteome Integral Solubility Alteration (5xMIC)
Proteome Integral Solubility Alteration (MIC)
HP_0914
HP_1512
(E)
Compound IVk
Proteome Integral Solubility Alteration (5xMIC)
Proteome Integral Solubility Alteration (MIC)
HP_1539
tupC

**Fig. S4 Concentration-dependent changes in Proteome Integral Solubility Alteration**

**(A)** Concentration-dependent targets for Compound IV: Shows the targets that exhibit a concentration-dependent response to Compound IV, represented by the teal color.
**(B)** Concentration-dependent targets for Compound IVa: Shows the targets that exhibit a concentration-dependent response to Compound IVa, represented by the teal color.
**(C)** Concentration-dependent targets for Compound IVb: Shows the targets that exhibit a concentration-dependent response to Compound IVb, represented by the teal color.
**(D)** Concentration-dependent targets for Compound IVj: Shows the targets that exhibit a concentration-dependent response to Compound IVj, represented by the teal color.
**(E)** We have not detected any significant concentration-dependent targets for Compound IVk.

(A)

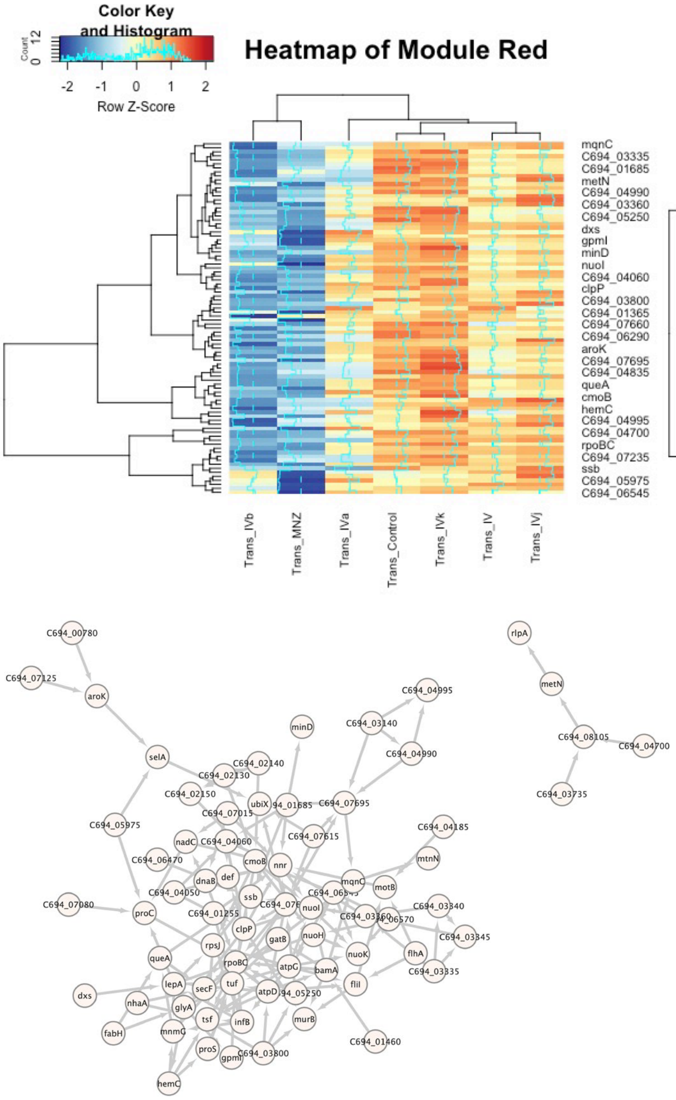


| Term ID | Term description | Strength | FDR |
|---|---|---|---|
| GO:0110165 | Cellular anatomical entity | 0.11 | 0.0084 |
| GO:0016020 | Membrane | 0.25 | 0.0118 |
| KW-1278 | Translocase | 0.99 | 0.0076 |

(B)

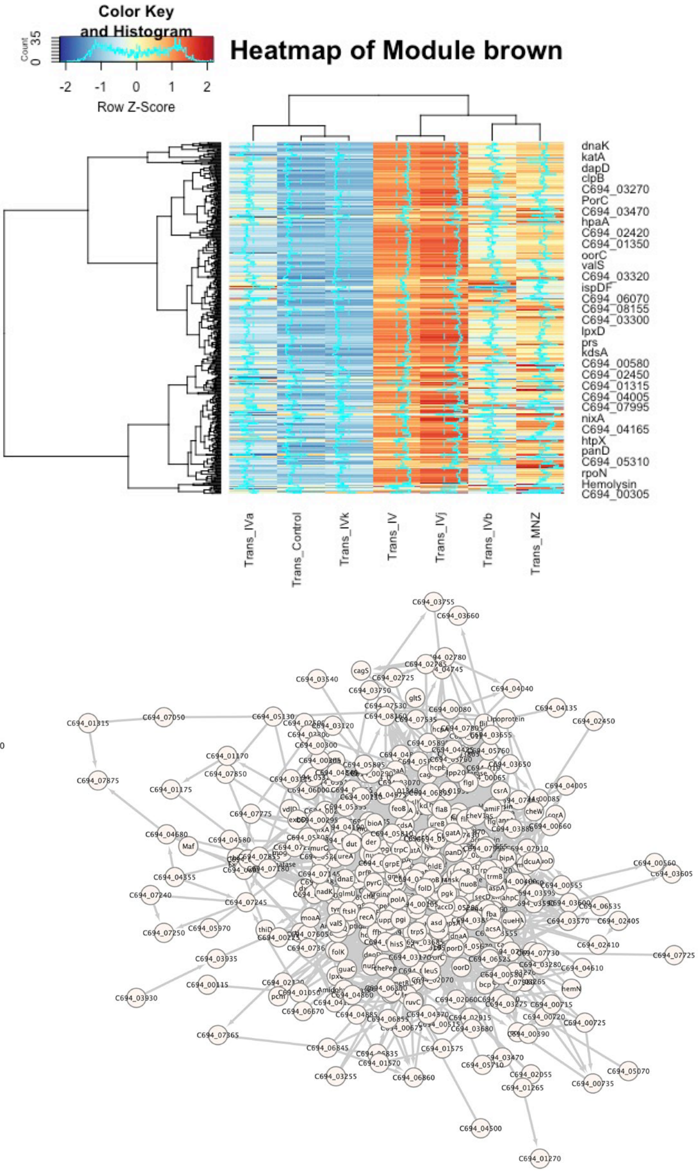


| Term ID | Term description | Strength | FDR |
|---|---|---|---|
| GO:0044281 | Small molecule metabolic process | 0.2 | 0.0060 |
| GO:0055114 | Oxidation-reduction process | 0.29 | 0.0060 |
| GO:0019752 | Carboxylic acid metabolic process | 0.25 | 0.0121 |
| GO:0003824 | Catalytic activity | 0.11 | 8.06e-05 |
| GO:0043167 | Ion binding | 0.14 | 0.0032 |
| GO:0016491 | Oxidoreductase activity | 0.28 | 0.0131 |
| GO:0036094 | Small molecule binding | 0.17 | 0.0156 |
| GO:0005737 | Cytoplasm | 0.13 | 0.00029 |
| GO:0005622 | Intracellular | 0.11 | 0.00088 |
| GO:0110165 | Cellular anatomical entity | 0.06 | 0.00088 |
| GO:0005829 | Cytosol | 0.14 | 0.0429 |
| CL:1197 | Carbon metabolism, and fatty acid metabolic process | 0.36 | 0.00068 |
| CL:1198 | Carbon metabolism, and fatty acid metabolic process | 0.4 | 0.00068 |
| CL:1199 | Carbon metabolism, and fatty acid synthase activity | 0.4 | 0.0026 |
| CL:1201 | Pyruvate metabolism, and Citrate cycle (TCA cycle) | 0.47 | 0.0330 |
| heo01100 | Metabolic pathways | 0.2 | 2.45e-05 |
| heo01110 | Biosynthesis of secondary metabolites | 0.26 | 0.00047 |
| heo01120 | Microbial metabolism in diverse environments | 0.31 | 0.0029 |
| heo01200 | Carbon metabolism | 0.38 | 0.0029 |
| heo02020 | Two-component system | 0.42 | 0.0101 |
| heo00020 | Citrate cycle (TCA cycle) | 0.49 | 0.0425 |
| heo01230 | Biosynthesis of amino acids | 0.28 | 0.0457 |

**Fig. S5 Gene expression heatmap and correlation of different compounds across modules from the WGCNA analysis.**

**(A)** Gene expression heatmap and correlation in module "red": This figure shows the gene expression patterns of Compound IV and its derivatives, along with their correlation in the "red" module. It also depicts the protein-protein coexpression network of the genes in this module and the associated gene ontology. **(B)** Gene expression heatmap and correlation in module "brown": This figure shows the gene expression patterns of Compound IV and its derivatives, along with their correlation in the "brown" module. It also illustrates the protein-protein coexpression network of the genes in this module and the associated gene ontology.

(A)

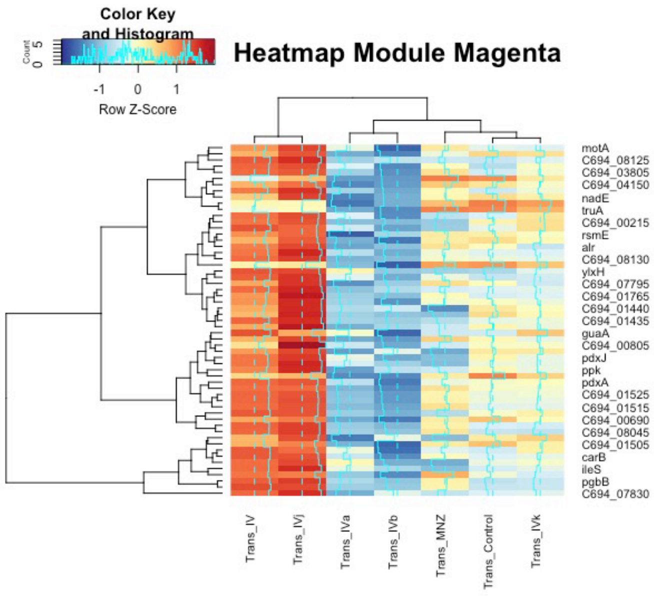


(B)

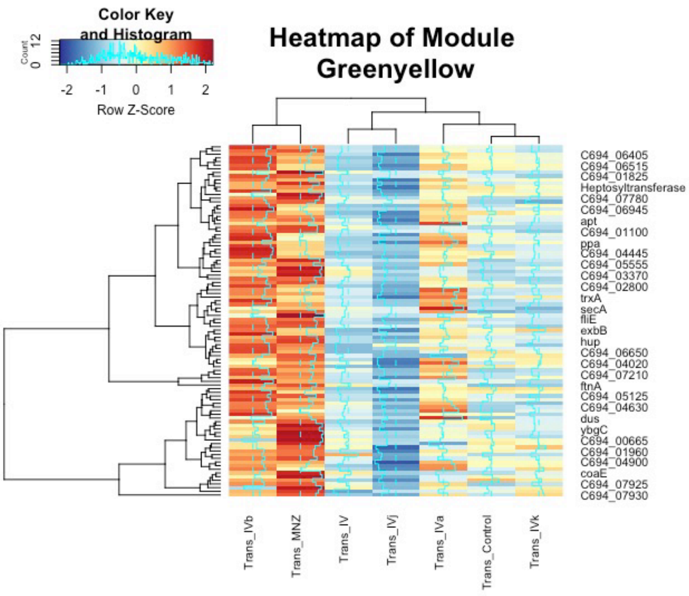


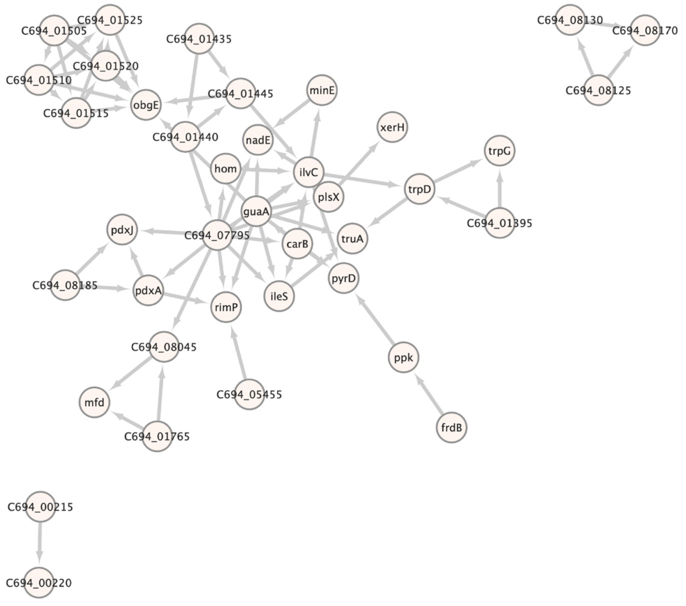


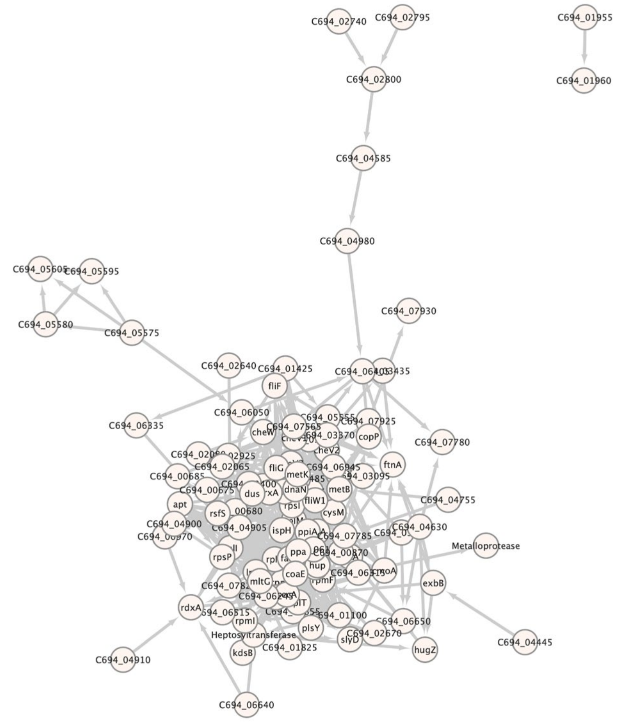


| Term ID | Term description | Strength | FDR |
|---|---|---|---|
| CL:2616 | Dipeptide transmembrane transporter activity, and Oligopeptide/dipeptide ABC transporter, C-terminal | 1.45 | 0.0033 |

| Term ID | Term description | Strength | FDR |
|---|---|---|---|
| GO:0006935 | Chemotaxis | 0.99 | 0.00013 |
| GO:0040011 | Locomotion | 0.78 | 0.00031 |
| GO:0007165 | Signal transduction | 0.88 | 0.0020 |
| GO:0042221 | Response to chemical | 0.68 | 0.0020 |
| GO:0007154 | Cell communication | 0.69 | 0.0108 |
| CL:1948 | Taxis | 1.1 | 9.53e-05 |
| CL:1869 | Locomotion, and cell projection organization | 0.58 | 0.0018 |
| CL:1870 | Locomotion, and cell projection organization | 0.59 | 0.0018 |
| CL:1871 | Locomotion, and cell projection organization | 0.61 | 0.0018 |
| CL:1951 | CheW-like domain, and Transducer | 1.14 | 0.0130 |
| heo02030 | Bacterial chemotaxis | 0.98 | 4.04e-05 |
| SM00260 | Two component signalling adaptor domain | 1.14 | 0.0109 |
| SM00283 | Methyl-accepting chemotaxis-like domains (chemotaxis sensory transducer). | 1.14 | 0.0245 |
| SM00448 | cheY-homologous receiver domain | 0.84 | 0.0327 |

**Fig. S6 Gene expression heatmap and correlation of different compounds across modules from the WGCNA analysis.**

**(A)** Gene expression heatmap and correlation in module “magents": This figure shows the gene expression patterns of Compound IV and its derivatives, along with their correlation in the " magents " module. It also depicts the protein-protein coexpression network of the genes in this module and the associated gene ontology. **(B)** Gene expression heatmap and correlation in module “Greenyellow": This figure shows the gene expression patterns of Compound IV and its derivatives, along with their correlation in the " Greenyellow " module. It also illustrates the protein-protein coexpression network of the genes in this module and the associated gene ontology.

(A)

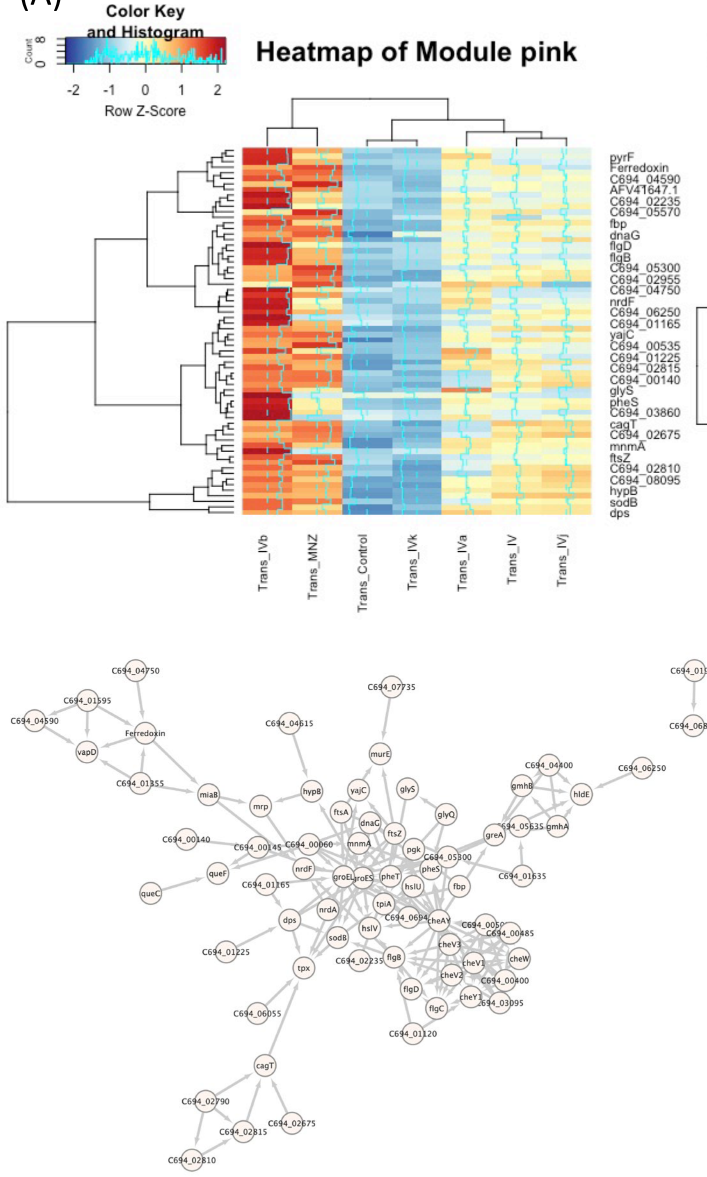


| Term ID | Term description | Strength | FDR |
|---|---|---|---|
| GO:0006935 | Chemotaxis | 1.04 | 0.00037 |
| GO:0007165 | Signal transduction | 1.01 | 0.00037 |
| GO:0040011 | Locomotion | 0.84 | 0.00037 |
| GO:0042221 | Response to chemical | 0.75 | 0.0012 |
| CL:1948 | Taxis | 1.23 | 5.89e-06 |
| CL:1871 | Locomotion, and cell projection organization | 0.67 | 0.0019 |
| CL:1951 | CheW-like domain, and Transducer | 1.27 | 0.0057 |
| CL:388 | Mixed, incl. protein folding, and detoxification | 0.68 | 0.0293 |
| CL:1097 | ADP-L-glycero-beta-D-manno-heptose metabolic process, and GmhA/DiaA | 1.17 | 0.0299 |
| CL:389 | Mixed, incl. protein folding, and detoxification | 0.7 | 0.0375 |
| heo02030 | Bacterial chemotaxis | 1.02 | 0.00015 |
| heo02020 | Two-component system | 0.72 | 0.0064 |
| PF01584 | CheW-like domain | 1.27 | 0.0234 |
| IPR002545 | CheW-like domain | 1.27 | 0.0369 |
| IPR036061 | CheW-like domain superfamily | 1.27 | 0.0369 |
| SM00260 | Two component signalling adaptor domain | 1.27 | 0.0028 |
| SM00283 | Methyl-accepting chemotaxis-like domains (chemotaxis sensory transducer). | 1.27 | 0.0082 |
| SM00448 | cheY-homologous receiver domain | 0.97 | 0.0090 |

(B)

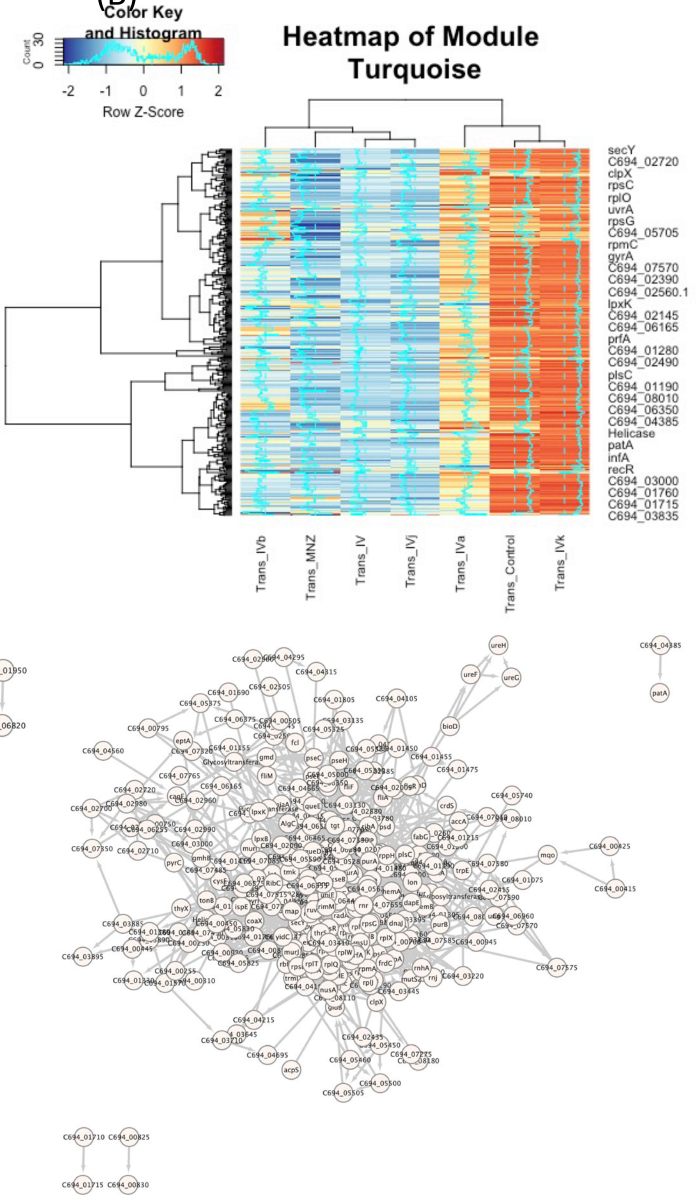


| Term ID | Term description | Strength | FDR |
|---|---|---|---|
| GO:0010467 | Gene expression | 0.32 | 6.00e-05 |
| GO:0009059 | Macromolecule biosynthetic process | 0.28 | 9.53e-05 |
| GO:0034645 | Cellular macromolecule biosynthetic process | 0.28 | 9.53e-05 |
| GO:0044238 | Primary metabolic process | 0.14 | 9.53e-05 |
| GO:0044249 | Cellular biosynthetic process | 0.19 | 9.53e-05 |
| GO:1901576 | Organic substance biosynthetic process | 0.19 | 9.53e-05 |
| GO:0071704 | Organic substance metabolic process | 0.12 | 0.00010 |
| GO:0043170 | Macromolecule metabolic process | 0.17 | 0.00014 |
| GO:0044271 | Cellular nitrogen compound biosynthetic process | 0.23 | 0.00022 |
| GO:0006412 | Translation | 0.35 | 0.00054 |
| GO:0043604 | Amide biosynthetic process | 0.32 | 0.00061 |
| GO:0008152 | Metabolic process | 0.1 | 0.00069 |
| GO:0044260 | Cellular macromolecule metabolic process | 0.18 | 0.00069 |
| GO:0006807 | Nitrogen compound metabolic process | 0.13 | 0.00070 |
| GO:0034641 | Cellular nitrogen compound metabolic process | 0.16 | 0.00070 |
| GO:0044237 | Cellular metabolic process | 0.11 | 0.00070 |
| GO:0019538 | Protein metabolic process | 0.25 | 0.0014 |
| GO:0044267 | Cellular protein metabolic process | 0.28 | 0.0016 |
| GO:0009987 | Cellular process | 0.07 | 0.0024 |

**Fig. S7 Gene expression heatmap and correlation of different compounds across modules from the WGCNA analysis.**

**(A)** Gene expression heatmap and correlation in module “pink": This figure shows the gene expression patterns of Compound IV and its derivatives, along with their correlation in the "pink" module. It also depicts the protein-protein coexpression network of the genes in this module and the associated gene ontology. **(B)** Gene expression heatmap and correlation in module “turquoise": This figure shows the gene expression patterns of Compound IV and its derivatives, along with their correlation in the " turquoise " module. It also illustrates the protein-protein coexpression network of the genes in this module and the associated gene ontology.

(A) (B)

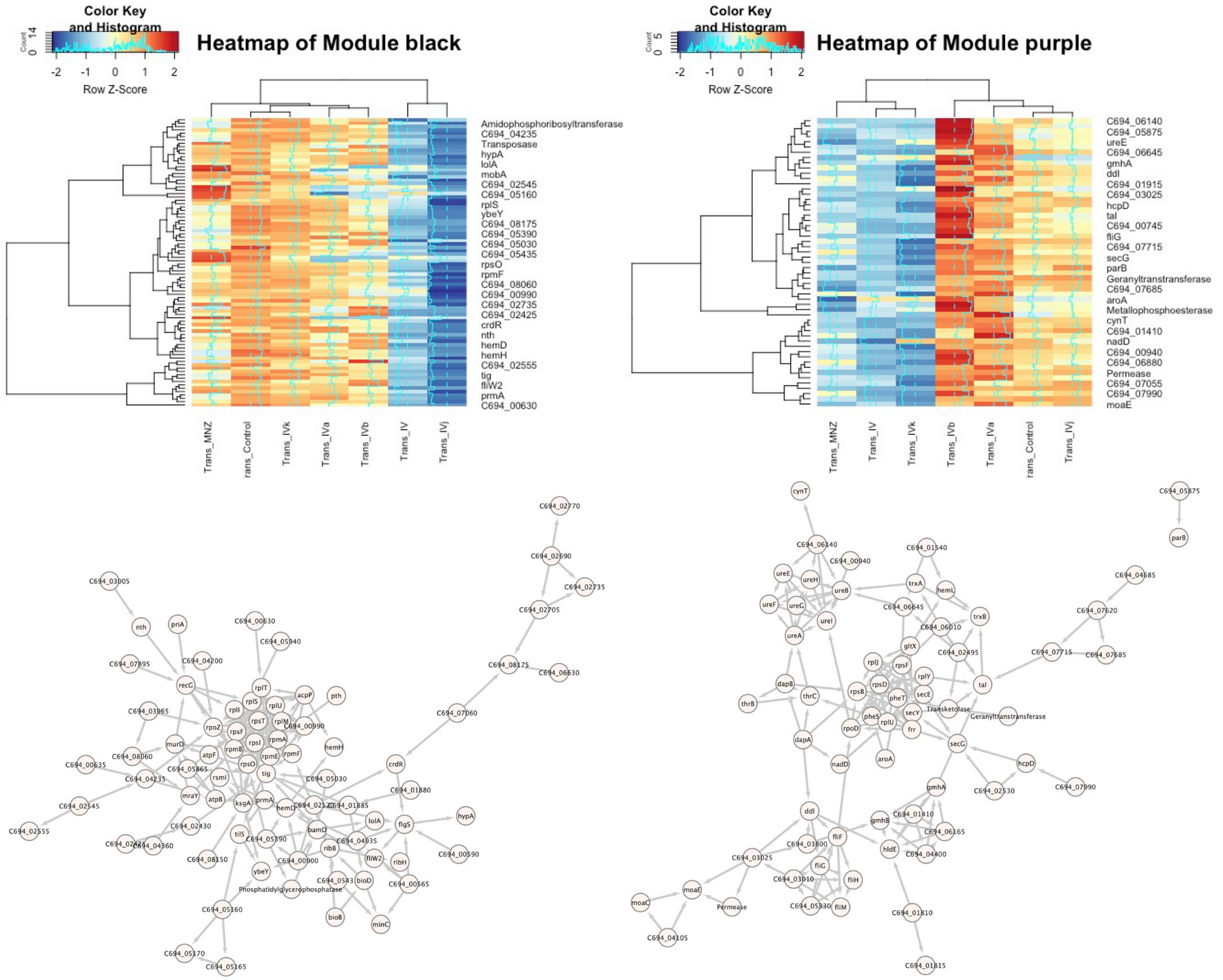


(A)

| Term ID | Term description | Strength | FDR |
|---|---|---|---|
| GO:0043170 | Macromolecule metabolic process | 0.25 | 0.0132 |
| GO:0044260 | Cellular macromolecule metabolic process | 0.27 | 0.0330 |
| GO:0003735 | Structural constituent of ribosome | 0.61 | 0.0396 |
| GO:0032991 | Protein-containing complex | 0.44 | 0.0018 |
| GO:0005840 | Ribosome | 0.59 | 0.0044 |
| GO:0043229 | Intracellular organelle | 0.5 | 0.0044 |
| GO:0043232 | Intracellular non-membrane-bounded organelle | 0.54 | 0.0044 |
| GO:1990904 | Ribonucleoprotein complex | 0.57 | 0.0100 |
| GO:0044391 | Ribosomal subunit | 0.58 | 0.0145 |
| GO:0022626 | Cytosolic ribosome | 0.59 | 0.0321 |
| CL:55 | Translation | 0.47 | 0.0203 |
| CL:57 | Ribosome, and transcription, DNA-templated | 0.52 | 0.0203 |
| CL:58 | Ribosome | 0.56 | 0.0203 |
| CL:63 | Ribosome | 0.6 | 0.0203 |
| CL:66 | Ribosome | 0.63 | 0.0203 |
| CL:51 | Translation, and RNA methylation | 0.39 | 0.0354 |
| CL:118 | Large ribosomal subunit | 0.79 | 0.0369 |
| CL:68 | Ribosome | 0.56 | 0.0498 |
| heo03010 | Ribosome | 0.61 | 0.0042 |
| KW-0689 | Ribosomal protein | 0.61 | 0.0096 |

(B)

| Term ID | Term description | Strength | FDR |
|---|---|---|---|
| GO:0009405 | Pathogenesis | 0.95 | 0.0201 |
| GO:0044419 | Interspecies interaction between organisms | 0.64 | 0.0299 |
| GO:0003774 | Motor activity | 1.15 | 0.0254 |
| GO:0016151 | Nickel cation binding | 1.05 | 0.0312 |
| GO:0110165 | Cellular anatomical entity | 0.12 | 0.0017 |
| GO:0005622 | Intracellular | 0.18 | 0.0178 |
| GO:0005737 | Cytoplasm | 0.18 | 0.0308 |
| CL:2821 | Nickel insertion, and urease activity | 1.27 | 0.0037 |
| CL:1877 | Motor activity, and Bacterial flagellum protein export | 0.97 | 0.0290 |
| CL:2796 | Nickel cation binding, and protein maturation | 0.82 | 0.0311 |
| KW-0843 | Virulence | 1.0 | 0.0061 |
| KW-0996 | Nickel insertion | 1.27 | 0.0371 |

**Fig. S8 Gene expression heatmap and correlation of different compounds across modules from the WGCNA analysis.**

**(A)** Gene expression heatmap and correlation in module “black": This figure shows the gene expression patterns of Compound IV and its derivatives, along with their correlation in the " black " module. It also depicts the protein-protein coexpression network of the genes in this module and the associated gene ontology. **(B)** Gene expression heatmap and correlation in module “purple": This figure shows the gene expression patterns of Compound IV and its derivatives, along with their correlation in the " purple" module. It also illustrates the protein-protein coexpression network of the genes in this module and the associated gene ontology.

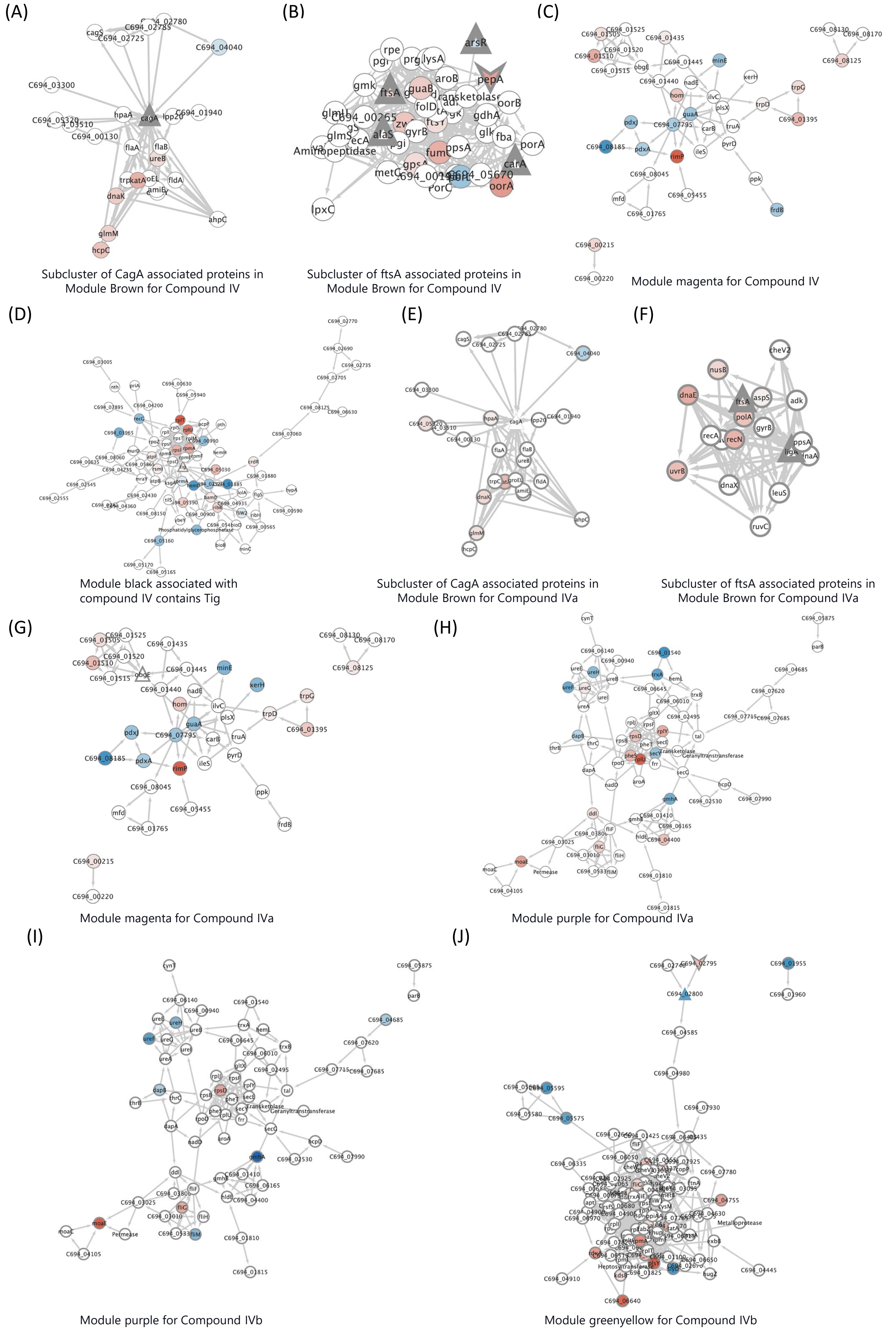
(A)
Subcluster of CagA associated proteins in Module Brown for Compound IV
(B)
Subcluster of ftsA associated proteins in Module Brown for Compound IV
(C)
Module magenta for Compound IV
(D)
Module black associated with compound IV contains Tig
(E)
Subcluster of CagA associated proteins in Module Brown for Compound IVa
(F)
Subcluster of ftsA associated proteins in Module Brown for Compound IVa
(G)
Module magenta for Compound IVa
(H)
Module purple for Compound IVa
(I)
Module purple for Compound IVb
(J)
Module greenyellow for Compound IVb

**Fig. S9 Associations between different modules and specific targets for Compound IV and its derivatives.**

**(A-B)** First neighbors of the target proteins CagA and FtsA from the "brown" module associated with Compound IV. **(C)** Module "magenta" associated with Compound IV. **(D)** Module "black" associated with Compound IV, with Tig as a target. **(E-F)** First neighbors of the target proteins CagA and FtsA from the "brown" module associated with Compound IVa. **(G)** Module "magenta" associated with Compound IVa, with ObgE as a target. **(H)** Module "purple" associated with Compound IVa. **(I)** Module "purple" associated with Compound IVb. **(J)** Module "greenyellow" associated with Compound IVb, with two uncharacterized proteins as targets.

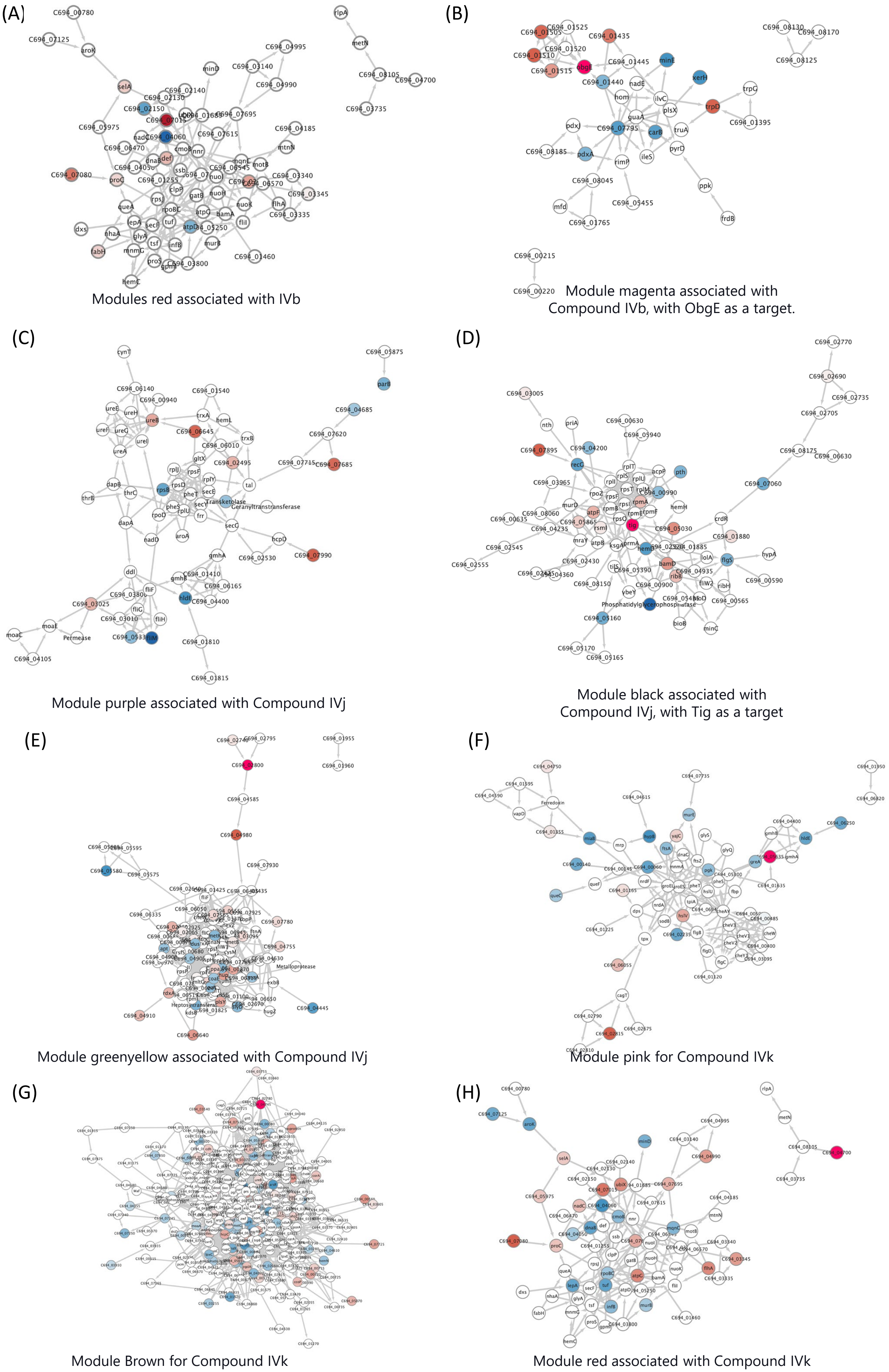
(A)
Modules red associated with IVb
(B)
Module magenta associated with Compound IVb, with ObgE as a target.
(C)
Module purple associated with Compound IVj
(D)
Module black associated with Compound IVj, with Tig as a target
(E)
Module greenyellow associated with Compound IVj
(F)
Module pink for Compound IVk
(G)
Module Brown for Compound IVk
(H)
Module red associated with Compound IVk

**Fig. S10 Associations between different modules and specific targets for Compound IV and its derivatives.**

**(A)** Module "red" associated with Compound IVb. **(B)** Module "magenta" associated with Compound IVb, with ObgE as a target. **(C)** Module "purple" associated with Compound IVj. **(D)** Module "black" associated with Compound IVj, with Tig as a target. **(E)** Module "greenyellow" associated with Compound IVj, with a hypothetical protein as a target. **(F)** Module "pink" associated with Compound IVk, with a hypothetical protein as a target. **(G)** Module "brown" associated with Compound IVk, with a hypothetical protein as a target. **(H)** Module "red" associated with Compound IVk, with a hypothetical protein as a target.

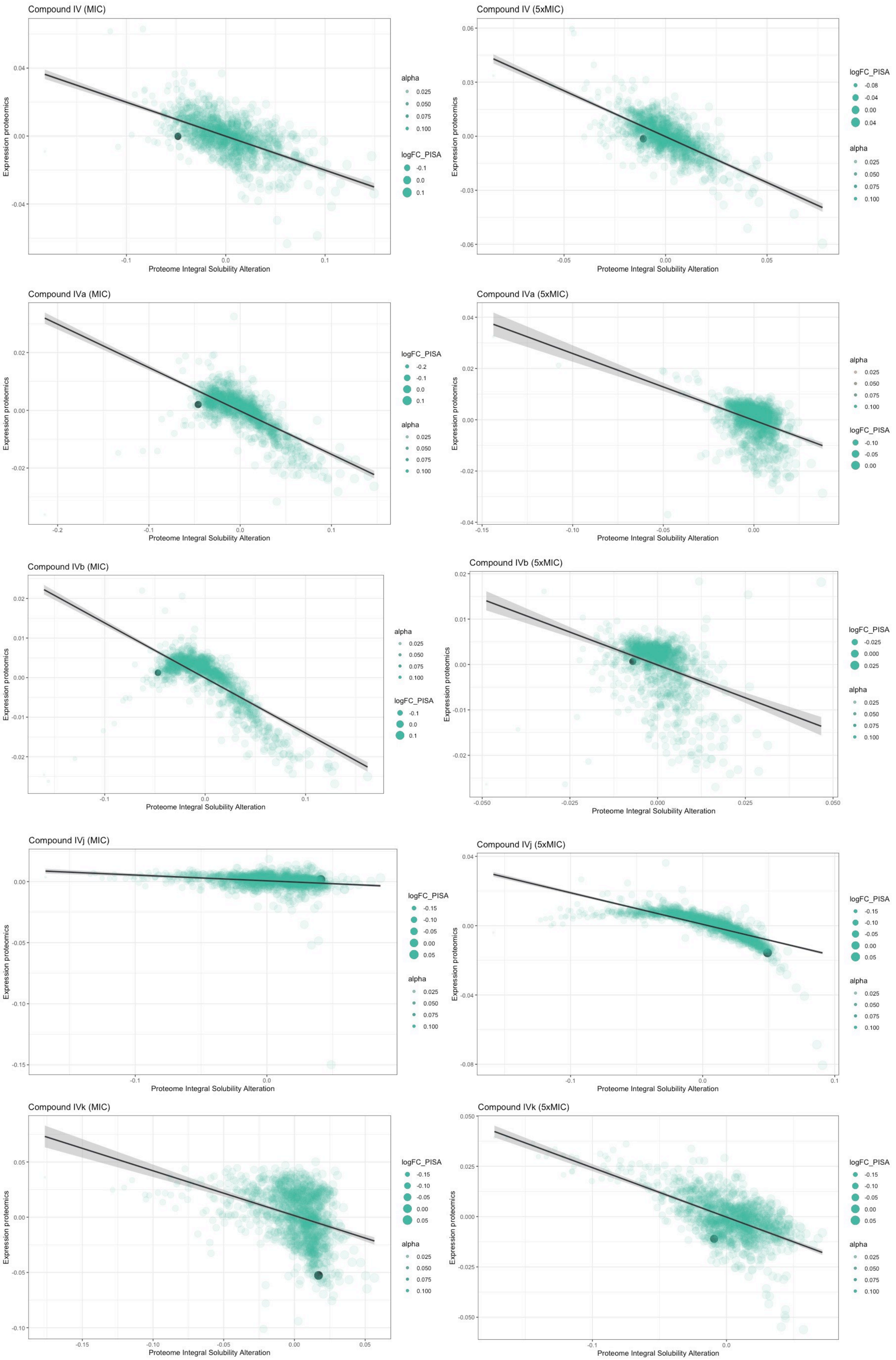
Compound IV (MIC)
Compound IV (5xMIC)
Compound IVa (MIC)
Compound IVa (5xMIC)
Compound IVb (MIC)
Compound IVb (5xMIC)
Compound IVj (MIC)
Compound IVj (5xMIC)
Compound IVk (MIC)
Compound IVk (5xMIC)
Expression proteomics
Proteome Integral Solubility Alteration
alpha
0.025
0.050
0.075
0.100
logFC_PISA

**Fig. S11 Alteration in solubility of flavodoxin in the Proteome Integral Solubility Alteration (PISA) assay associated with different compounds at two different concentrations.**

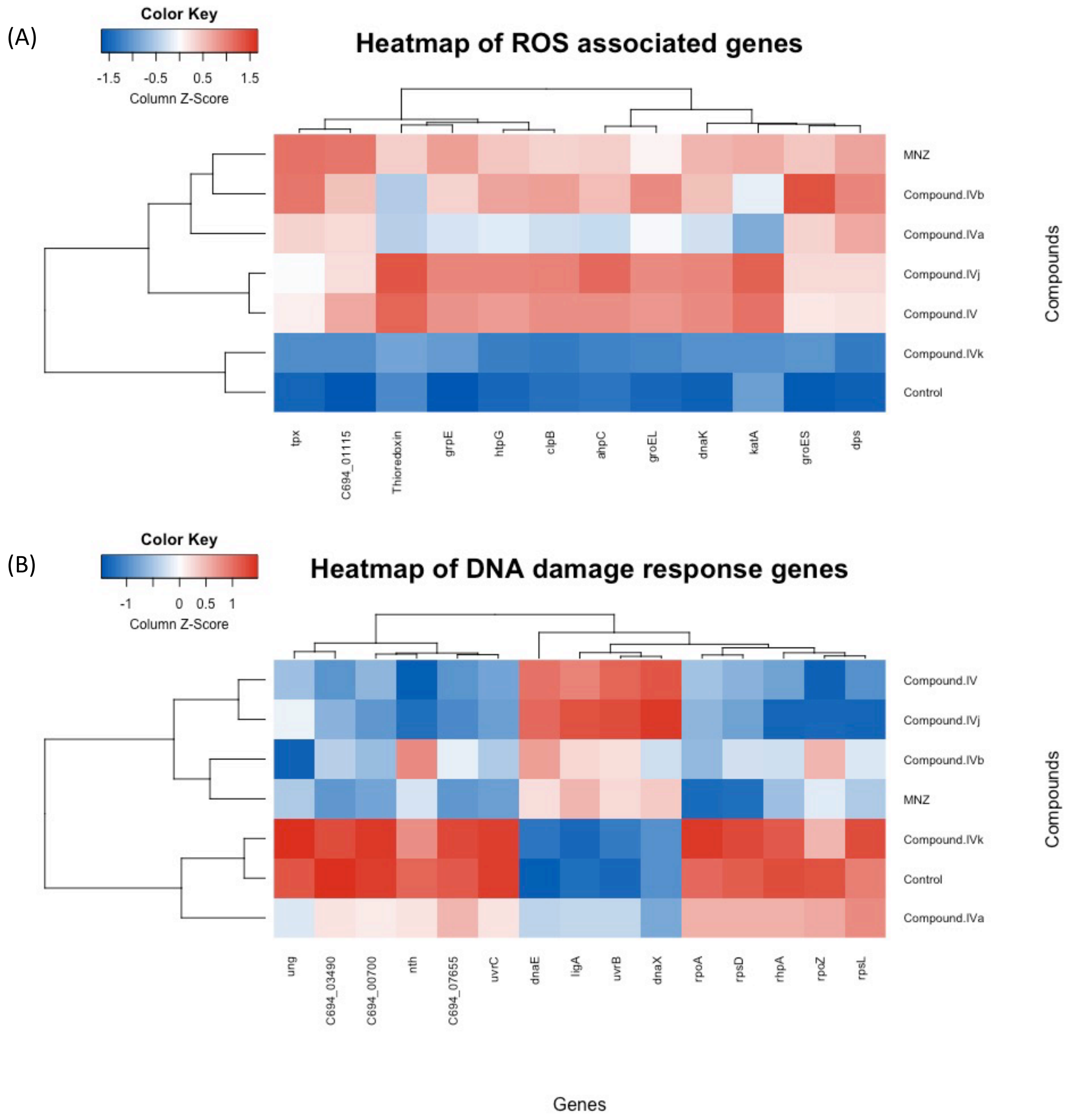


**Fig. S12 Heatmap of the differential gene expression associated with (A) generation of ROS and (B) DNA damage response genes.**
